# Highly efficient induction of functionally mature excitatory neurons from feeder-free human ES/iPS cells

**DOI:** 10.1101/2023.02.10.528087

**Authors:** Zhi Zhou, Wataru Kakegawa, Koki Fujimori, Misato Sho, Rieko Shimamura, Sopak Supakul, Sho Yoshimatsu, Jun Kohyama, Michisuke Yuzaki, Hideyuki Okano

## Abstract

Cortical excitatory neurons (Cx neurons) are the most dominant neuronal cell type in the cerebral cortex, which play a central role in cognition, perception, intellectual behavior and emotional processing. Robust *in vitro* induction of Cx neurons may facilitate as a tool for the elucidation of brain development and pathomechanism of the intractable neurodevelopmental and neurodegenerative disorders including Alzheimer’s disease, and thus potentially contribute to drug development. Here, we report a defined method for efficient induction of Cx neurons from the feeder-free-conditioned human embryonic stem cells (ES cells) and induced pluripotent stem cells (iPS cells). By using this method, human ES/iPS cells could be differentiated into ~99% MAP2-positive neurons by three weeks, and these induced neurons, within five weeks, presented various characteristics of mature excitatory neurons such as strong expression of glutamatergic neuron-specific markers (subunits of AMPA and NDMA receptors and CAMKIIα), highly synchronized spontaneous firing and excitatory postsynaptic current (EPSC). Moreover, the Cx neurons showed susceptibility to the toxicity of Aβ_42_ oligomers and excitotoxicity of excessive glutamates, which is another advantage in terms of toxicity test and searching for the therapeutic agents. Taken together, this study provides a novel research platform for the study of neural development and degeneration based on the feeder-free human ES/iPS cell system.

## Introduction

Cortical excitatory neurons (Cx neurons) account for approximately 80% of all neuronal cells in the cerebral cortex, in which they play a central role in cognition, perception, intellectual behavior and emotional processing [1]. Using glutamate as their neurotransmitter, Cx neurons, also known as projection neurons, are incorporated in high-order neuronal networks, and their activity is locally modulated by inhibitory GABAergic neurons representing approximately 20% of neuronal population within the cerebral cortex. Due to their important roles, the developmental abnormalities and the degeneration of Cx neurons could result in a variety of intractable neurological diseases, for examples, autism spectrum disorder (ASD), frontotemporal dementia (FTD), and Alzheimer’s disease (AD). Moreover, previous studies demonstrated that concentrating neurons with a region-specific identity made it possible to recapitulate disease-associated *in vitro* phenotypes [2–4]. Thus, a robust induction of Cx neurons *in vitro* holds a great promise for high-throughput screening of therapeutic candidates for these diseases.

Human embryonic stem cells (ES cells) and induced pluripotent stem cells (iPS cells) have potentials for unlimited proliferation and differentiation into all the three germ layers [5,6]. Based on pioneer studies on neural differentiation of mouse and human ES/iPS cells [7,8], induction of Cx neurons from human ES/iPS cells have been reported from various groups [2,9–16]. However, there remain several technical and biological limitations for these methods. For example, first, the reported methods used on-feeder human ES/iPS cells as a starting material, in which the maintenance of on-feeder human ES/iPS cells requires heterogenous feeder cells such as primary mouse embryonic fibroblasts (MEFs) or STO feeder cells [17]. Feeder cell preparation including a large-scale expansion and a mitotic inactivation necessitates significant effort. Moreover, due to the difficulty in completely elimination of the feeder cells, remaining feeders may hamper successful differentiation of a directed linage(s). Second, several reported methods utilized 3D culture system. Direct floating culture of ES/iPS cells, so called embryoid body (EB) [18,19], neurosphere [2] or organoid [14,15], usually results in a population of differentiated cells with developmental heterogeneity (neural stem cells located in the center, and neurons in the edge). Moreover, the cell-autonomous formation of anterior-posterior and dorsal-ventral axes in floating cell aggregates during 3D culture is frequently observed. Importantly, these 3D methods necessitate a long-term differentiation culture (over 2~3 months) in order to obtain functional mature neurons, which may be costly and not favorable for the high-throughput screening. To circumvent these limitations, we here report a defined method for the direct differentiation of Cx neurons from feeder-free human ES/iPS cells. By the step-by-step optimization of the differentiation procedures, we demonstrated a successful induction of Cx neurons with high efficiency (~ 100%).

## Material and Methods

### Ethical statements

Experiments using human ES/iPS cells were approved by the Ministry of Education, Culture, Sports, Science, and Technology (MEXT) of Japan and Ethics Committee of Keio University School of Medicine (approval number: 20080016). Recombinant DNA experiments were approved by the Recombinant DNA Experiment Safety Committee of Keio University.

### Cell culture

We used two human iPS cell lines (201B7 and 1210B2) and a human ES cell line (KhES1) in the present study, which were kindly provided by Drs. Shinya Yamanaka and Norio Nakatsuji (Kyoto University) [6,20,21]. We also used a disease-specific iPS cell line PS1 (A246E) established from a familial Alzheimer’s disease patient harboring an Ala246Glu mutation in *Presenilin 1* [22]. These ES/iPS cells were maintained in a feeder-free condition as described previously [20,23,24]. In brief, the ES/iPS cells were maintained in StemFit/AK02N (Wako (Ajinomoto), AJ100). We used 0.5× TrypLE select (Thermo Fisher Scientific) in PBS(-) for passaging and single-cell dissociation, and plated onto a tissue culture 6-well plate adding the AK02N medium supplemented with 10 μM Y27632 (Wako, 253-00513) for initial 24 h. The culture plates were coated with 1.5 μg/ml iMatrix-511 silk (Laminin-511 E8; Wako (Nippi), 381-07363) beforehand. Medium change was performed every other day, and passaging was performed every 7-10 d.

### Neuronal differentiation

For neuronal differentiation, single-cell-dissociated human ES/iPS cells were plated onto a Laminin-511 E8-coated tissue culture plate (4.17 × 10^4^ cells/cm^2^, 4.0 × 10^5^/well for 6-well plates, 1.62 × 10^5^/well for 12-well plates, 7.83 × 10^4^/well for 24-well plates, 3.1 × 10^4^/well for 48-well plates) in the AK02N medium supplemented with 10 μM Y27632 (Y27632 was only added for the initial 24 h). After plating, the ES/iPS cells were initially maintained in AK02N medium until the cell density has become sub-confluent (80 – 90%). Then, the medium was changed to a GMEM/KSR medium supplemented with 2 μM DMH1 (Wako, 041-33881), 2 μM SB431542 (Sigma, S4317) and 10 μM Y27632 (defined as 0 day(s) *in vitro*; 0 div). The GMEM/KSR medium was consisted of 1 × Glasgow’s MEM (GMEM; Wako, 078-05525) supplemented with 8% KnockOut Serum Replacement (KSR; Thermo Fisher Scientific, 10828-028). In the initial optimization experiments, we also used MHM/B27 [2] and N2/B27 media [25]. From 1 to 14 div, medium change was performed every day using the same medium, which Y27632 was withdrawn from 1 div, and DMH1 and SB431542 was withdrawn from 8 div. On 15 div, cells were washed in PBS once, and then incubated with 1 × Accutase (Nacalai Tesque, 12679-54) for 30 m at 37°C for single-cell dissociation. The dissociated cells were initially collected in GMEM/8%KSR with 10 μM Y27632. After centrifuging the cell suspension (200 × g, 5 m) and removing the supernatant, cells were suspended in a Plating medium. The Plating medium was consisted of 1 × BrainPhys Neuronal Medium/N2-A/SM1 kit (BrainPhys; Stem Cell Technologies, 05793) supplemented with 10 μM Y27632, 10 ng/ml BDNF (R&D Systems, 248-BD-025), 10 ng/ml GDNF (Peprotech, 450-10), 200 μM Ascorbic acid (AA; Sigma, A4403-100MG), 0.5 mM dbcAMP (Sigma, D0627-1G), 10 μM DAPT (Sigma, D5942-5MG) and 2 μM PD0332991 (Sigma, PZ0199). Prior to plating, we filtrated the cell suspension using a 40-μm cell strainer to remove aggregated cells. The cells were plated onto a Poly-L-Lysine and Laminin-coated 24-well plate at a density of 4.0 x 10^5^ cells/well. Poly-L-Lysine and Laminin coating of 24-well plates was performed by 48 h incubation at 37°C using Tissue Culture water (Sigma, W3500) supplemented with 1 μg/ml Poly-L-Lysine (Sigma, P4832) and 4 ng/ml Mouse Laminin (Thermo Fisher Scientific, 23017015). On 18 div, half medium change was performed using a Maintenance medium with 10 μM DAPT. The Maintenance medium was consisted of 1 × BrainPhys supplemented with 10 ng/ml BDNF, 10 ng/ml GDNF, 200 μM Ascorbic acid, 0.5 mM dbcAMP and 2 μM PD0332991. On 21 div, full medium change was performed using the Maintenance medium. From 24 div, medium change was performed every 3 d using the same medium.

### qRT-PCR

RNA extraction, reverse transcription, and qRT-PCR was performed as described previously [26]. The level of *ACTB* expression was used as an internal control for normalization. As a control of cerebral cortex, we used total cDNA obtained from an adult human sample.

### Immunocytochemistry

Immunocytochemistry was performed as described previously [26]. For high-content quantitative immunocytochemical analysis, we used IN Cell Analyzer 6000 (GE Healthcare) as described previously [3]. The detailed analysis protocol using IN Cell Analyzer 6000 is available upon request. Information of primary and secondary antibodies used in this study are available from the corresponding authors.

### ELISA

Enzyme-linked immunosorbent assay (ELISA) was performed as described previously [27]. In brief, we collected culture media of Cx neurons. The collected media were briefly centrifuged to remove insoluble material and can be kept at −80°C until the analysis. The remaining cells were lysed in RIPA buffer (Wako) and protein concentration was measured by BCA Protein assay (Pierce). A *β* 40 and Aβ42 levels in the media were measured using commercial kits, Human βAmyloid (1–40) ELISA Kit II (Wako, 298–64601) and Human βAmyloid (1–42) ELISA Kit High Sensitive (Wako, 296–64401) according to the manufacturer’s introductions. Each Aβ concentration was normalized by protein levels of the culture cells.

### Electrophysiology and Ca imaging

Microelectrode array (MEA) recording was performed using the Maestro system (Axion Biosystems) as described previously [28]. In brief, 12-well MEA plates were pre-coated using Poly-L-Lysine and Laminin, and neural progenitors (on 15 div) were subsequently plated onto the electrode area in the MEA plate. Data were acquired using a sampling rate of 12.5 kHz and filtered using a 200–3000 Hz Butterworth band-pass filter. Detection threshold was set to +6.0 × SD of the baseline electrode noise. Spike raster plots were analyzed using Neural Metric Tool (Axion Biosystems). The spike count files generated from the recordings were used to calculate the number of active electrodes (defined as an electrode which has an average of more than 5 spikes/min) in each well, the average per-active electrode mean firing rate (MFR or spikes/min) and the standard deviation of the average per-active electrode MFR. The data from the initial 3 min in each data file were omitted to enable the activity to stabilize in the Maestro, and 10–15 min of activity was subsequently recorded.

Calcium (Ca) imaging was performed as described previously [23]. In brief, Fluo-8 (AAT Bioquest; final 5 μM), Probenecid (final 1 mM), Pluonic F-127 (final 0.02%) and Hoechst (final 0.5 μg/ml) were supplemented into the plating medium and incubated for 30 min at 37°C. After incubation, the cells were washed twice with PBS and fed with fresh medium. For TTX or CNQX treatment, the reagent was supplemented with a final concentration of 2 μM or 50 μM, respectively. Movies were captured using an IX83 inverted microscope (Olympus) equipped with an electron multiplying CCD camera (Hamamatsu Photonics) and pE-4000 LED illumination system (CoolLED). We recorded the Ca oscillation of the cells using the MetaMorph Image Analysis Software in following parameters: 12 outputs, 750 msec interval for 2 min, GFP filter, 1x gain, 2.75 MHz, and 80 msec exposure. Regions of interest (ROIs) were drawn on cells based on time projection images of the recordings. ROI traces of the time course of changes in green fluorescence intensity were generated and used as substrates for subsequent analyses. To adjust for photobleaching, the difference in intensity between the first and last frames was calculated and subtracted from the raw intensity. The change in fluorescence intensity over time was normalized as ΔF/F= (F-F0)/F0, where F0 is fluorescence at the starting point of exposure (time 0). ΔFmax was defined as the difference of the largest change in ΔF/F. Whole-cell patch clump was performed as described previously [26].

### Statistical Analysis

All data were expressed as means ± SD. Statistical significance of differences was analyzed with the Welch’s *t*-test. Differences of *P* < 0.05 were expressed as *, *P* < 0.01 as **, and *P* < 0.001 were as ***, which were considered statistically significant.

## Results

### Efficient induction of neural progenitors with a cerebrocortical identity from feeder-free human iPS cells

Initially, we sought to devise an optimized induction method of SOX1(+) incipient ectodermal cells from 201B7 healthy-control human iPS cells [6] cultured in a feeder-free condition using AK02N as a culture media and iMatrix-511 as a coating materials [20,24]. Based on the principle methodology described in previous studies [2,9–13,29,30], we testified a 7-days culture of iPS cells using three media, such as N2B27 [25], MHM/B27 [2] and GMEM/KSR [31] without supplementation of any chemical compounds (see Material and Methods). FACS analysis using cell-permeable antibodies revealed that, among the culture media, GMEM/KSR medium resulted in the highest yield of SOX1(+) cells, compared to those of N2B27 and MHM/B27 media (Supplementary Figure 1A).

Next, we performed simultaneous quantification of SOX1(+) cells and OCT4(+) pluripotent cells since the remaining pluripotent cells may hamper the subsequent neuronal differentiation due to their high proliferation rate and unpredictable differentiation into other lineages. In addition, we tested supplementation of a BMP4 inhibitor, DMH1, and TGF-β inhibitor, SB431542, to prevent the cells from the non-ectodermal differentiation [7]. When using the N2B27 medium, although DMH1 and SB431542 treatment ameliorated the SOX1(+)/OCT4(-) rates, we observed approximately 30–60% remaining OCT4(+) cells (Supplementary Figure 1B). On the other hand, when using the GMEM/KSR medium, we found higher rates of SOX1(+)/OCT4(-) cells and very low rates of OCT4(+) cells (~ 5%) in the combination with DMH1 and SB431542 treatment (Supplementary Figure 1C). Therefore, we decided to focus on the method using the GMEM/KSR medium supplemented with DMH1 and SB431542 during the initial phase of induction (Left part of Figure 1A) for the further analyses. Using total RNA derived from the differentiated cells at 12 div (days *in vitro*) and the undifferentiated iPS cells, we compared the gene expressions of pluripotency (*NANOG* and *POU5F1*), neuroectoderm (*SOX1* and *PAX6*) and cerebral cortex (*FOXG1* and *SIX*3)-related marker genes (Figure 1B). As a result, the differentiated cells showed significant upregulation of *SOX1, PAX6*, *FOXG1*, and *SIX3*, and downregulation of *NANOG* and *POU5F1* compared to the iPS cells (Figure 1B). Since low-dose treatment (each 2 μM) of DMH1 and SB431542 showed a marginal increase in *SOX1* expression compared to that of high-dose treatment (5 and 10 μM) (data not shown), subsequent experiments were performed by the low-dose condition. In the condition, we also assessed time-dependent change of the marker genes’ expression by qRT-PCR (Supplementary Figure 2A) and immunocytochemistry (Supplementary Figure 2B). Collectively, these data demonstrated that our optimized method enabled a robust induction of neural progenitors with an identity of cerebral cortex.

**Figure 1.**
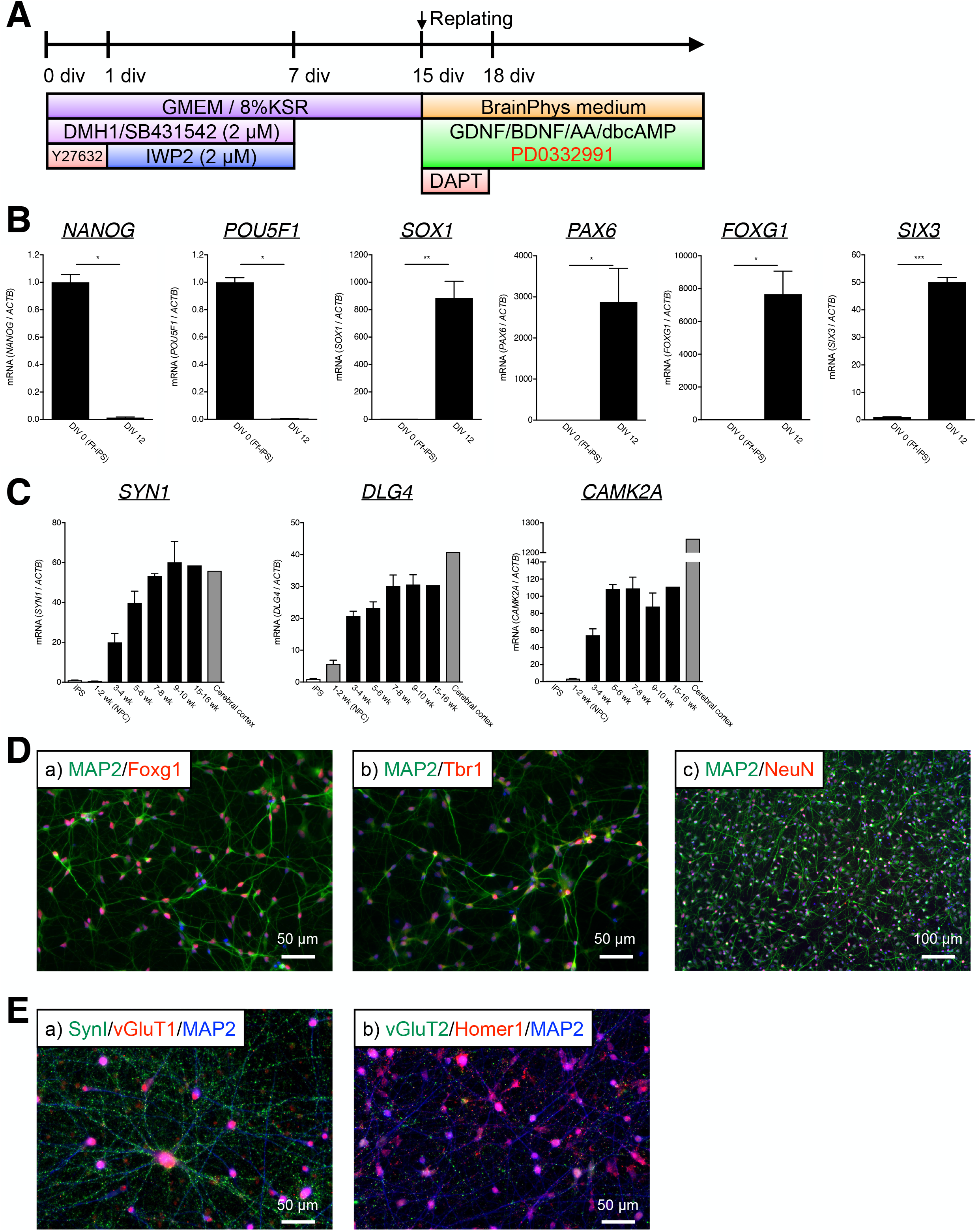
Optimization of the induction method. (A) Graphical timetable of the induction method. For the detail, see the “Neuronal differentiation” section in Material and Methods. (B) qRT-PCR analysis of pluripotency (*NANOG* and *POU5F1*), neuroectoderm (*SOX1* and *PAX6*) and cerebral cortex (*FOXG1* and *SIX*3)-related markers in the original feeder-free-conditioned 201B7 iPS cells (DIV 0 Ff-iPS) and the differentiated cells on div 12 (DIV 12). (C) qRT-PCR analysis of synaptic markers (*SYN1* and *DLG4*) and *CAMK2A*. (D) Immunocytochemical analysis of the differentiated cells (5 weeks after differentiation). Nuclei were stained blue using Hoechst. (E) Immunocytochemical analysis of the Cx neurons (10 weeks after differentiation).

### Maturation culture of neural progenitors with a CDK inhibitor

By single-cell dissociation and subsequent replating of neural progenitors on 15 div, we next attempted further differentiation of these cells into functional mature neurons (Right part of Figure 1A). Based on our preliminary results that the BrainPhys-based medium, not Neurobasal-based medium, enhanced neuronal survival and spontaneous firing activity of the differentiated cells evaluated by Ca imaging (data not shown), we utilized the BrainPhys medium supplemented with GDNF, BDNF, ascorbic acid (AA) and dbcAMP for neuronal survival and maturation. We also supplemented a γ-secretase inhibitor DAPT and CDK4/6 inhibitor PD0332991 for further neuronal differentiation through the promotion of cell cycle exit [16]. Since prolonged γ-secretase inhibition may impair neuronal function [32], DAPT was withdrawn on 21 div. For the ease of further analyses, we initially compared three coating conditions before replating of neuronal progenitors as follows: Poly-L-Lysine/Laminin, Matrigel and Laminin only. Six days after replating, we found the least number of cell aggregations in the Poly-L-Lysine/Laminin condition (Supplementary Figure 3A). Using the optimized BrainPhys medium and coating condition, we observed a time-dependent increase of *SYN1, DLG4 (PSD-95*) and *CAMK2A* gene expression upon 5–6 or 7–8 weeks after differentiation of iPS cells (Figure 1C). We also confirmed the increasing of glutamate ionotropic receptor genes essential for functional maturation of glutamatergic neurons (Supplementary Figure 3B), and Alzheimer’s disease-associated marker Tau (*MAPT*) and its postnatal-specific 4R isoform (Supplementary Figure 3C). In addition, immunocytochemical analysis revealed that most of the differentiated cells (5 weeks after differentiation) were positive for MAP2, Foxg1 and NeuN (Figure 1D). Given these results, the differentiated cells for 5 weeks or more are hereinafter referred to as the Cx neurons. Moreover, we discovered that the Cx neurons (10 weeks after differentiation) showed dot-like immunoreactivity of synaptic markers including SynI, vGluT1, vGluT2 and Homer1, implying the formation of functional synapses (Figure 1E).

### Electrophysiological analyses of the Cx neurons

To ascertain the formation of functional synapses and local circuits, we performed Calcium (Ca) imaging analysis of the Cx neurons using a fluorescent Ca indicator Fluo-8 (Material and Methods). We recorded Ca dynamics on 15+7, 15+21, 15+36 and 15+94 div (Video S1–4). While differentiated cells on 15+7 and 15+21 div showed only spontaneous firing-like Ca flux (Video S1–2), Cx neurons on 15+36 and 15+94 div showed synchronized firing-like Ca flux (Video S3–4). Moreover, using the Cx neurons on 15+36 div, we demonstrated that the treatment with a sodium channel blocker TTX induced the complete loss of spontaneous firing-like Ca flux (data not shown), and the treatment with an AMPA/kainate receptor antagonist CNXQ resulted in no synchronization of Ca flux (Figure 2A), indicating that these synchronized Ca dynamics may be caused by functional synaptic activity.

**Figure 2.**
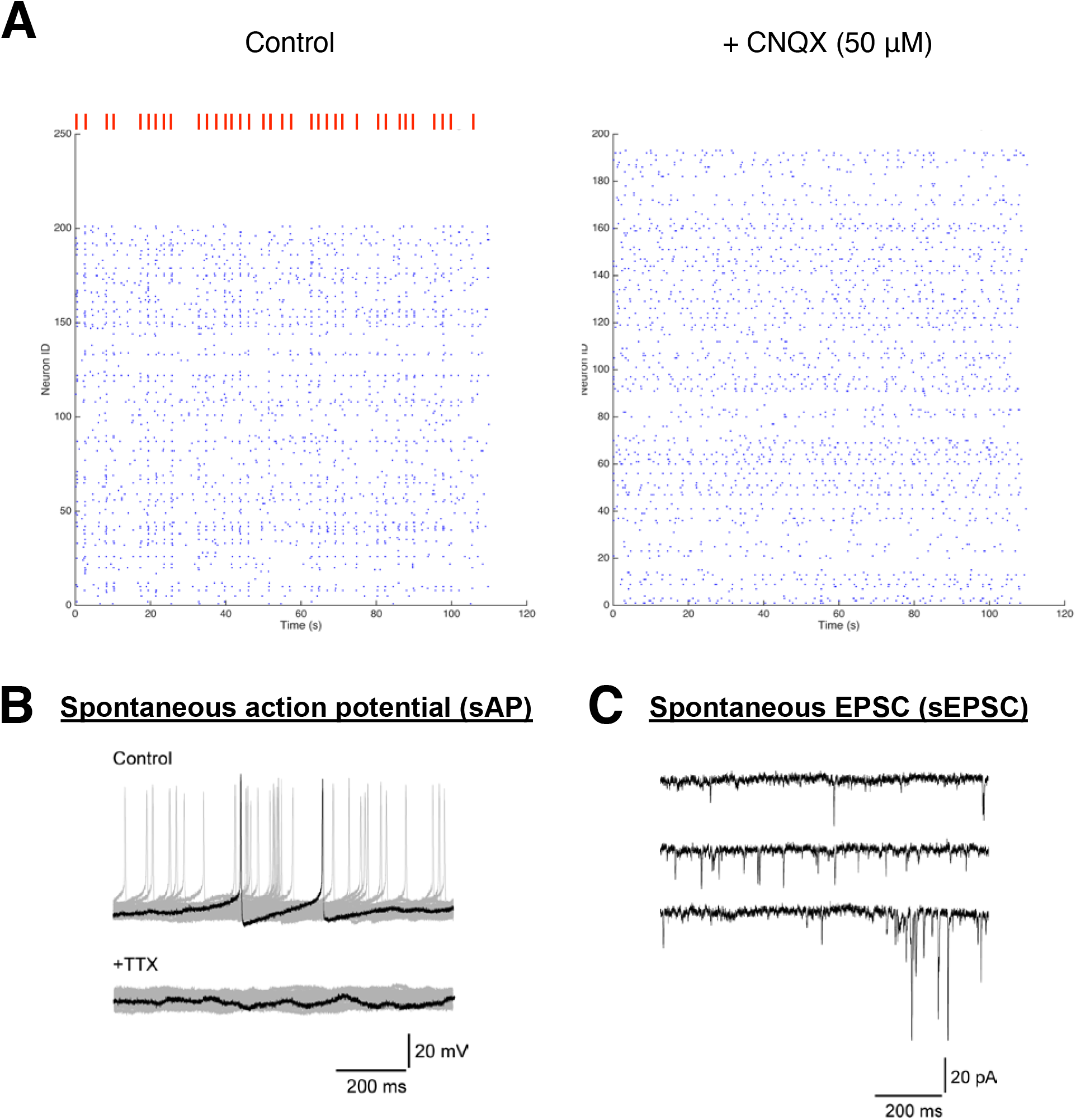
Electrophysiological analysis. (A) Ca imaging analysis using Fluo-8 in the Cx neurons on 15+36 div (Movie S3). Spikes were counted based on ΔF/F of the Fluo-8 fluorescence intensity. The vertical axis shows Neuron ID, and horizontal axis shows time (sec). Synchronized spike, only found in control cells, are depicted as red lines. (B–C) Whole-cell patch clump of the Cx neurons on 15+34 div.

Furthermore, to directly assess the electrophysiological characteristics of the Cx neurons, we carried out a whole-cell patch clamp and microelectrode array (MEA) analysis (Figure 2B–C and Supplementary Figure 4). The whole-cell patch clump showed the spontaneous action potential (sAP) the Cx neurons (15+34 div; 10 out of 16 cells) (Figure 2B, top) and excitatory postsynaptic current (sEPSC) (Figure 2C). The sAP was subsequently dissipated after the TTX treatment (Figure 2B, bottom). By MEA analysis, we detected synchronized spontaneous firing of the Cx neurons on 15+35 div (Supplementary Figure 4). Taken together, we demonstrated that the Cx neurons were functional mature, and able to form local neuronal circuits through the excitatory synapses.

### Susceptibility to toxic agents

To explore the glutamatergic neuron-specific susceptibility of the Cx neurons to the known neurotoxic agents, we initially tested the supplementation of excessive L-glutamate (L-Glu) inducing cell death by excessive excitotoxicity through glutamate receptors [33]. As shown in Figure 3A, we performed high-content imaging analysis of the Cx neurons using IN CELL Analyzer. L-Glu toxicity was evaluated by MAP2(+) neurite length and WST-8 bioreduction based cell viability assay. As a result, supplementation of excessive glutamate, 12.5 *μ*M or more, caused a decrease in neurite length per cell (Figure 3A–B) and WST-8 bioreduction activity (Figure 3C). On the other hand, supplementation of an NMDA glutamate receptor antagonist MK801 (1 *μ* M) partially attenuated the toxicity of excessive L-Glu (Figure 3A–C), indicating that the effect was caused by glutamate receptors which are expressed in the Cx neurons.

**Figure 3.**
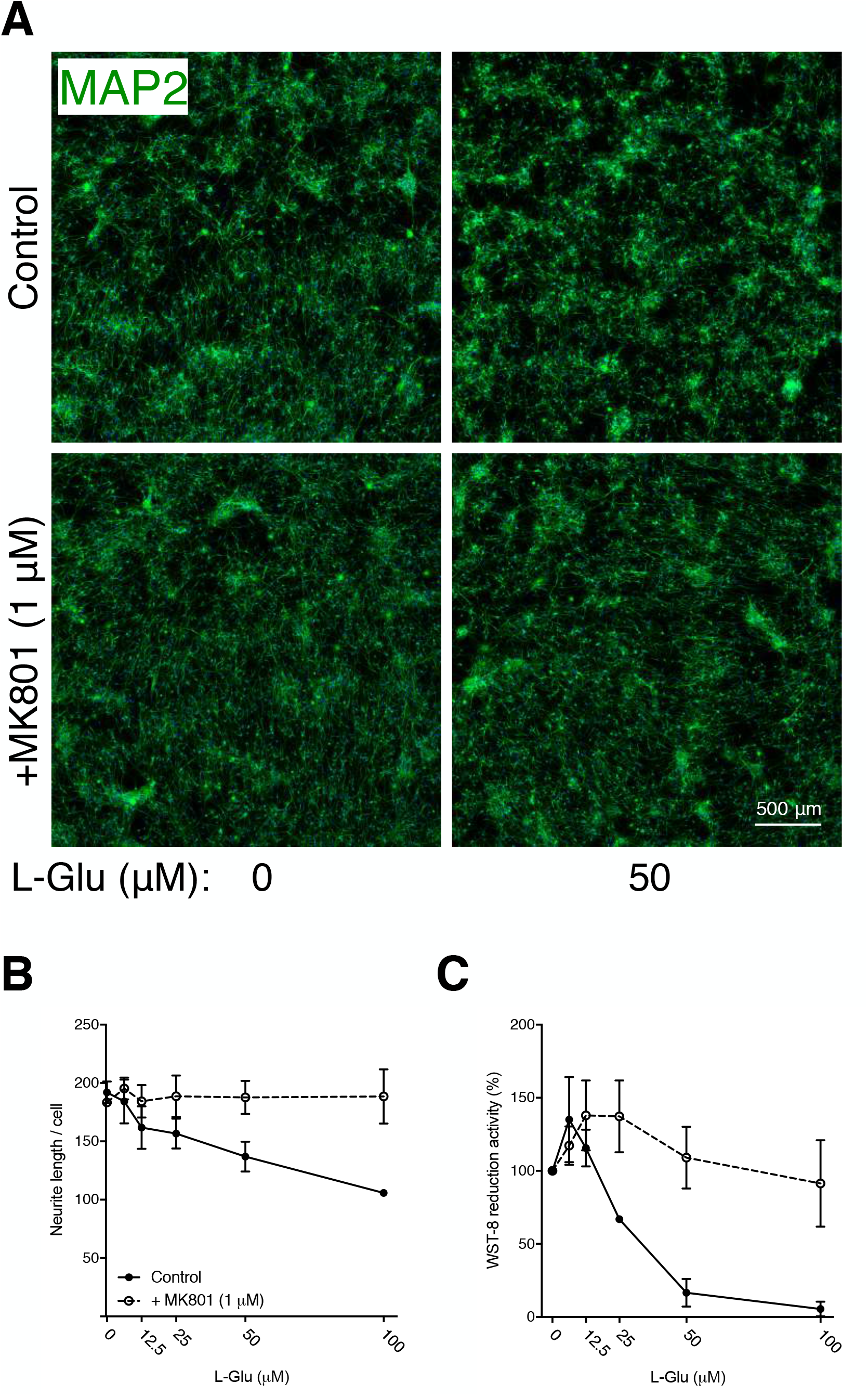
Susceptibility of the Cx neurons to L-Glu. (A) Representative immunocytochemical images of the Cx neurons with/without MK801 (1 μM) and L-Glu (50 μM) supplementation. Neurites were stained green with the MAP2 antibody. (B–C) Dose-dependent toxicity of L-Glu (0, 12.5, 25, 50 and 100 μM) with/without MK801, analyzed by MAP2(+) neurite length (B) and WST-8 bioreduction activity (C). The absorbance at 460 nm of the Cx neurons (0 μM L-Glu, MK801(-) condition) is set to 100%.

Next, we addressed the applicability of the Cx neurons for the *in vitro* modeling of Alzheimer’s disease. Supplementation of A *β* 42 oligomer in the Cx neuron culture (Figure 4A) resulted in a dose-dependent decrease in neurite length per cell (Figure 4B) and WST-8 bioreduction activity (Figure 4C). Furthermore, we assessed endogenous secretion of A *β* 40 and Aβ42 from the Cx neurons. Supplementation of DAPT dramatically decreased the secretion of Aβ40 (Figure 4D). Using the Cx neurons differentiated from a familial Alzheimer’s patient-derived iPS cell line, PS1 (A246E) [22], we found that disease-specific phenotypes such as decreased Aβ40 secretion (Figure 4E, left), increased Aβ42 secretion (Figure 4E, center) and thus increased ratio of Aβ42/Aβ40 (Figure 4E, right) were recapitulated. Thus, we demonstrated that utilizing the Cx neurons were advantageous for recapitulating region-specific neurological pathology.

**Figure 4.**
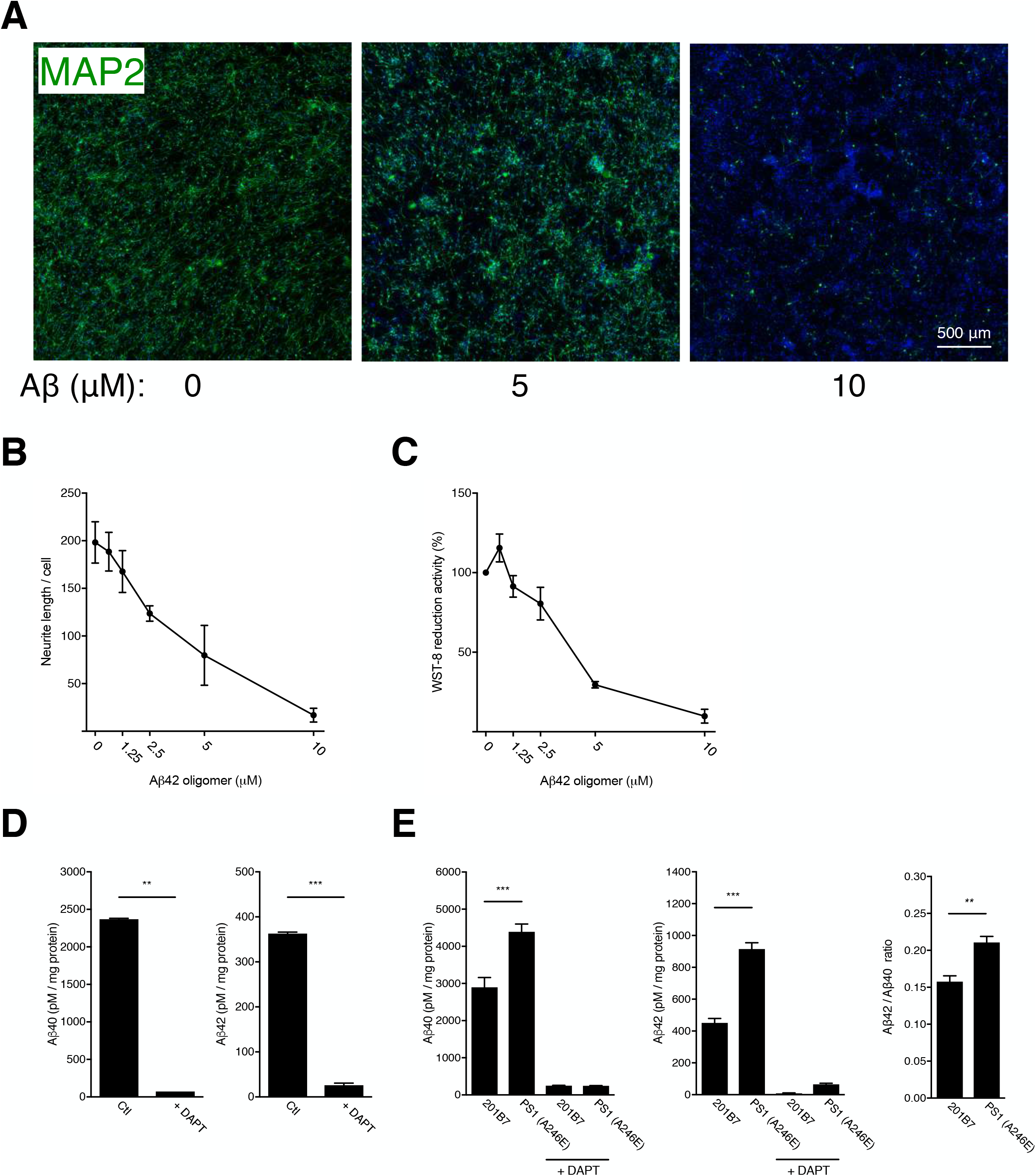
Susceptibility of the Cx neurons to Aβ oligomer and endogenous Aβ secretion. (A) Representative immunocytochemical images of the Cx neurons (201B7) with/without A *β* 42 oligomer (0, 5, 10 μM). Neurites were stained green with the MAP2 antibody. (B–C) Dose-dependent toxicity of Aβ42 oligomer (0, 1.25, 2.5, 5 and 10 μM), analyzed by MAP2(+) neurite length (B) and WST-8 bioreduction activity (C). The absorbance at 460 nm of the Cx neurons (0 μM Aβ42 oligomer condition) is set to 100%. (D) ELISA of endogenous Aβ40 and Aβ42 secretion from the Cx neurons (differentiated from 201B7 and PS1 (A246E) iPS cells).

### Reproducibility of the induction method in multiple human ES/iPS cell lines

Lastly, we assessed the reproducibility of the induction method using other human ES/iPS cell lines with different origins, such as 1210B2 and KhES1 (Supplementary Figure 5A). We showed that these two lines were successfully differentiated into MAP2(+) positive neurons at high efficiencies comparable to that of 201B7 (data not shown). Moreover, using the electrophysiology analysis, we demonstrated that the Cx neurons derived from these lines exhibited a maturity which was susceptible to the toxicity test of L-Glu (Supplementary Figure 5B) and Aβ42 oligomer (Supplementary Figure 5C).

## Discussion

In the present study, we report the defined induction method of Cx neurons from the feeder-free human ES/iPS cells. We highlight three advantages of this method as following: (i) High efficiency of neuronal induction with a region (cerebral cortex)-specific identity, which is important for recapitulation of disease-specific phenotypes [2–4]. (ii) High reproducibility in multiple cell lines, which is essential for unbiased comparison and robust applicability for disease-specific lines. In particular, since a major part of underlying genetic/epigenetic mechanisms of sporadic diseases remains unclear, we believe that *in vitro* systems with high purity and reproducibility could tackle this issue. (iii) Ease of the method. Our initial motivation was to explore the ease-of-use for high-throughput screening using human ES/iPS cells. To this end, we exploited feeder-free human ES/iPS cells, and also, succeeded in shortening the induction period required for obtaining functional mature neurons (upon 5 weeks). This makes it possible to perform cost-effective screening of therapeutic candidates using Cx neurons.

Amyloid beta, including Aβ40 and Aβ42. is the major components of senile plaques found in the brain of Alzheimer’s disease patients. Whether and how amyloid beta causes neurodegeneration in patients’ brain are yet unclear and under debate, our data suggested that Cx neurons induced by our method could be directly applicable for the elucidation of this mechanism *in vitro* as the endogenous Aβ secretion activity was enough for recapitulating the mutant Presenilin 1-associated Alzheimer’s disease phenotype (Figure 4D–E). However, while adding DAPT, the γ-secretase inhibitor, could possibly alter the results from the analysis of Aβ, our additional inductions of other cell lines, which were not originally included in this study, have pointed out the importance of adding DAPT to yield a high quality Cx neurons since cells demonstrated more survival rates with the supplementation of DAPT (described in a subsequent study).

Furthermore, functional maturation of the Cx neurons, presented by our data in electrophysiology analysis (Figure 2 and Supplementary Figures 4 and 5C), *CAMK2A* and 4R-Tau expression (Figure 1C and Supplementary Figure 3D), also indicated that the induced Cx neurons are also applicable for recapitulation of the late-onset disease phenotypes. Although further analyses are still required, we demonstrated the reported method here may serve as a novel research platform for deciphering cerebral cortex-specific pathomechanisms of neurological disorders, and thus might also contribute to drug screening and development.

## Acknowledgement

We especially thank Drs. Hirotaka Watanabe and Mitsuru Ishikawa (Keio University) for helpful discussion, and all the laboratory members of H.O. and M.Y. for encouragement and generous support for this study. This study was funded by the MEXT and the AMED (Grant ID: JP20dm0207001 to H.O.).

