## Supplementary material for "Highly efficient induction of functionally mature excitatory neurons from feeder-free human ES/iPS cells": hiPS-Cx_supplementary_2022bioRxiv.docx

**Supplementary Figure Legends**

**Supplementary Figure 1. Optimization of the SOX1(+)/OCT4(-) cell induction.**

(A−C) FACS analysis of the differentiated cells on ~ div (A) and ~ div (B−C) using cell-permeable Alexa647-conjugated Oct4 (Oct3/4) and PE-conjugated Sox1 antibodies.

**Supplementary Figure 2. Optimization of the induction of neural progenitors with a cerebral cortex identity.**

(A) qRT-PCR analysis of pluripotency (*NANOG* and *POU5F1*), neuroectoderm (*SOX1* and *PAX6*), cerebral cortex (*FOXG1*) and neuronal (*TUBB3*)-related marker genes.

(B) Representative immunocytochemical images of differentiated cells on div 0, 1, 3, 5, 7, 9 and 11.

**Supplementary Figure 3. Neuronal maturation culture.**

(A) Representative phase-contrast images of the differentiated cells (15 + 6 div) at low (top) and high (bottom) magnification.

(B−C) qRT-PCR analysis of the original 201B7 iPS cells, cells differentiated for 1-2, 3-4, 5-6, 7-8, 9-10 and 15-18 weeks, and control human cerebral cortex using specific primers of glutamate receptor genes.

(D) qRT-PCR analysis of total tau (left) and its 4R isoform (right).

**Supplementary Figure 4. MEA analysis.**

Using the Cx neurons on 15+35 div, we measured voltage change recorded from each electrode (left). From the data, spike rates (spike / sec) were measured (right).

**Supplementary Figure 5. Reproducibility of the method.**

(A) Summary of human ES/iPS cell lines used in this study.

(B) Dose-dependent toxicity of L-Glu (0, 12.5, 25, 50 and 100 μM) with/without MK801, analyzed by MAP2(+) neurite length (top) and WST-8 bioreduction activity (bottom) using the Cx neurons from 1210B2 (left) and KhES1 (right) lines. The absorbance at 460 nm of the Cx neurons (0 μM L-Glu, MK801(-) condition) is set to 100%.

(E) Dose-dependent toxicity of Aβ42 oligomer (0, 1.25, 2.5, 5 and 10 μM), analyzed by MAP2(+) neurite length (top) and WST-8 bioreduction activity (bottom) using the Cx neurons from 1210B2 (left) and KhES1 (right) lines. The absorbance at 460 nm of the Cx neurons (0 μM Aβ42 oligomer condition) is set to 100%
