## Supplementary material for "Highly efficient induction of functionally mature excitatory neurons from feeder-free human ES/iPS cells": Sup Fig5.pdf

# A

| Line: | 201B7 | 1210B2 | KhES1 |
|---|---|---|---|
| **Origin** | Adult human fibroblast | Adult peripheral blood mononuclear cells (PBMC) | human embryo |
| **Reprogramming** | Retrovirus (OCT3/4, SOX2, KLF4, c-MYC) | Episomal vectors (OCT3/4, SOX2, KLF4, L-MYC, dominant negative p53, EBNA1) | - |

# B

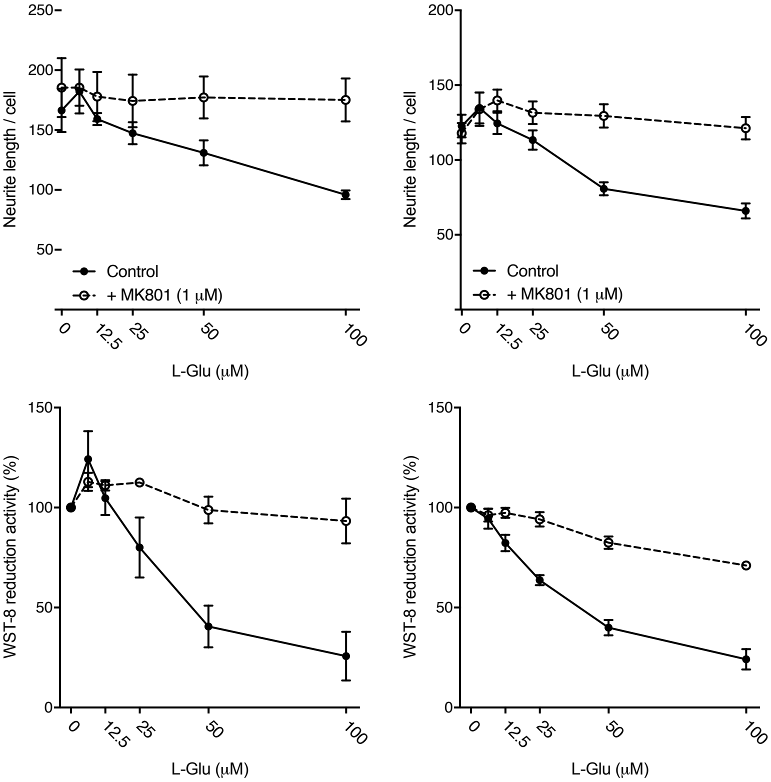


# C

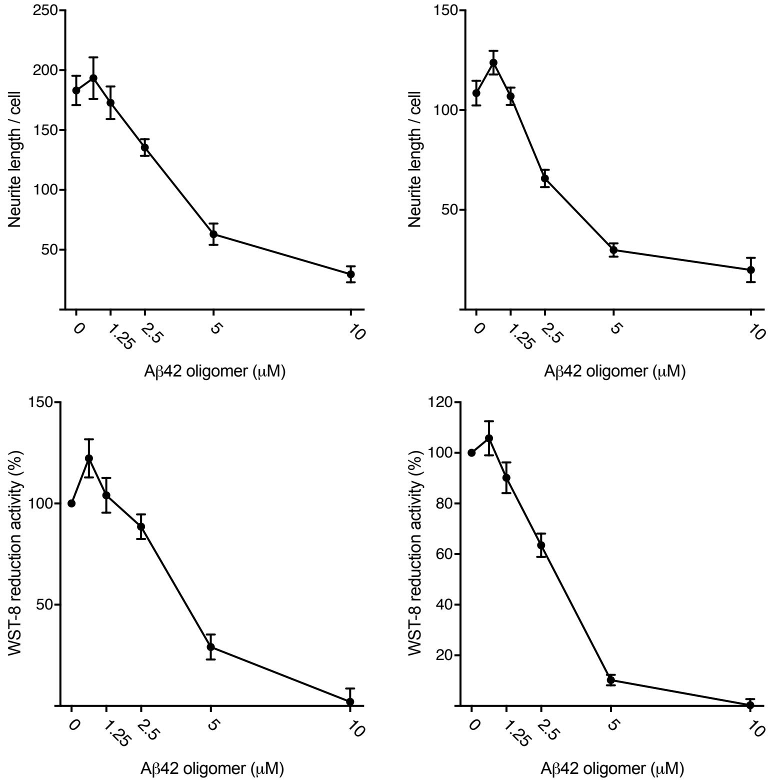


**Figure. S5 (Zhou *et al.*)**
