## Supplementary figures and images for "Highly efficient induction of functionally mature excitatory neurons from feeder-free human ES/iPS cells"

### Sup Fig1.pdf

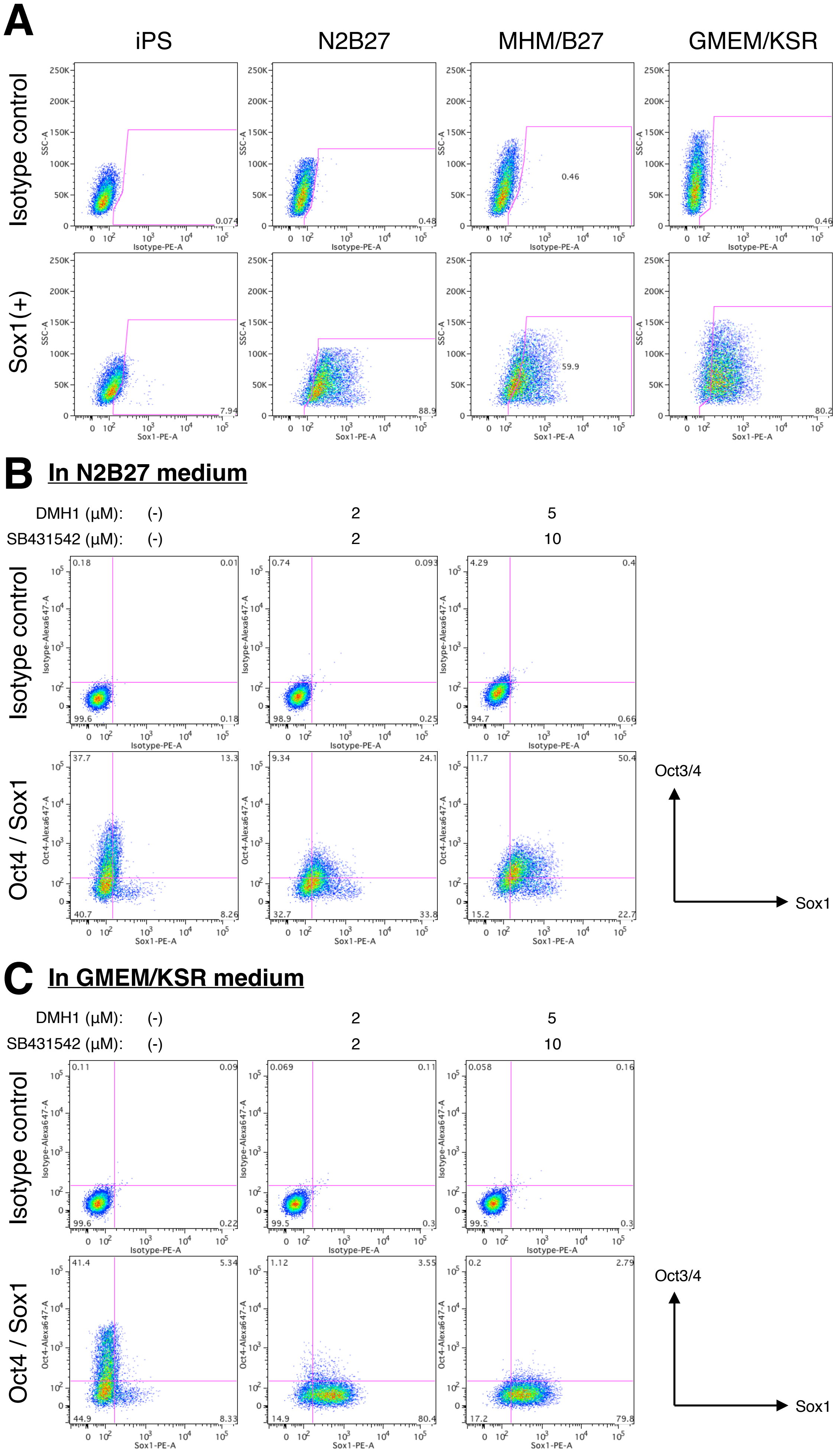


**Figure. S1 (Zhou *et al.*)**

### Sup Fig2.pdf

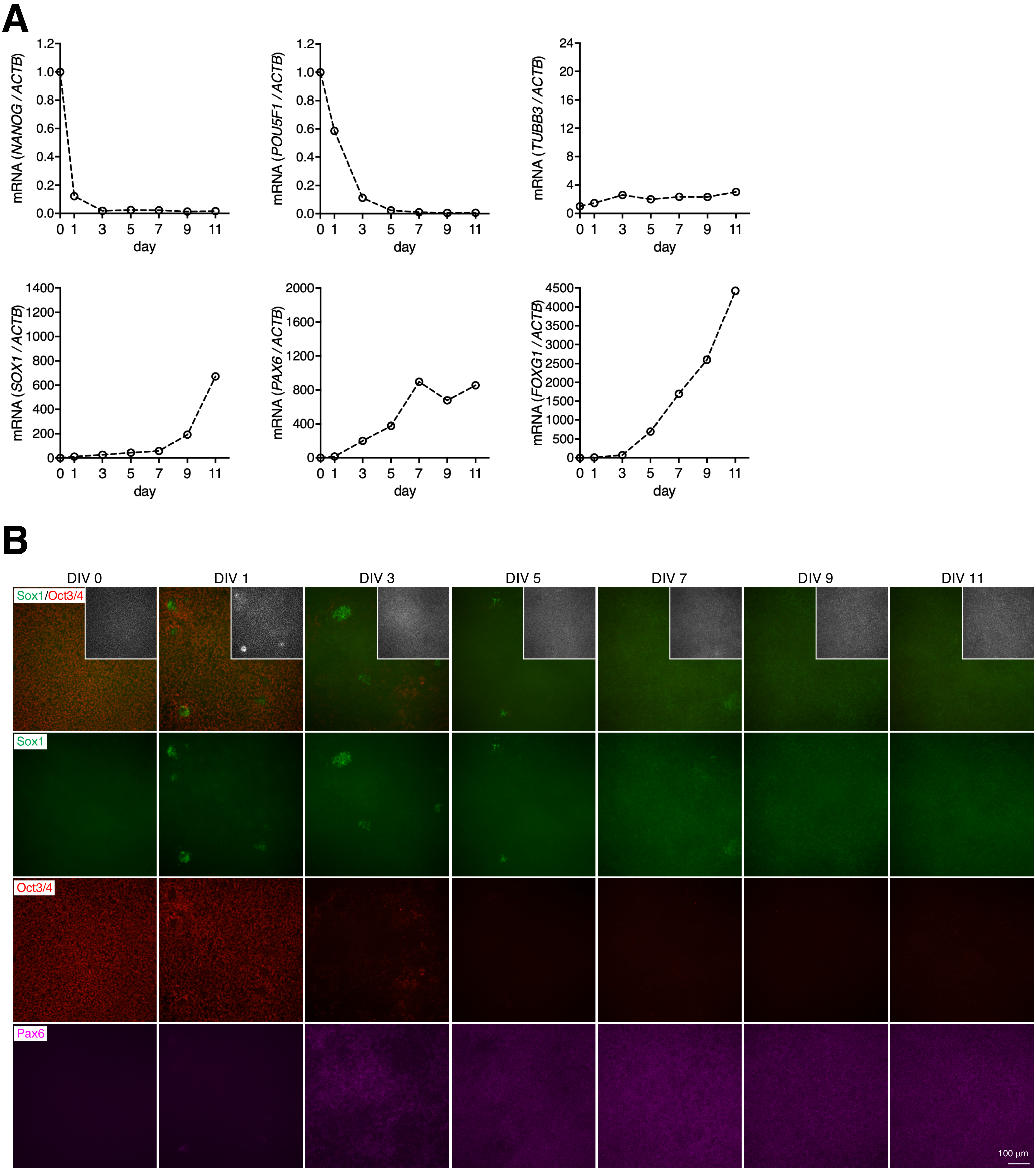


Figure. S2 (Zhou *et al.*)

### Sup Fig3.pdf

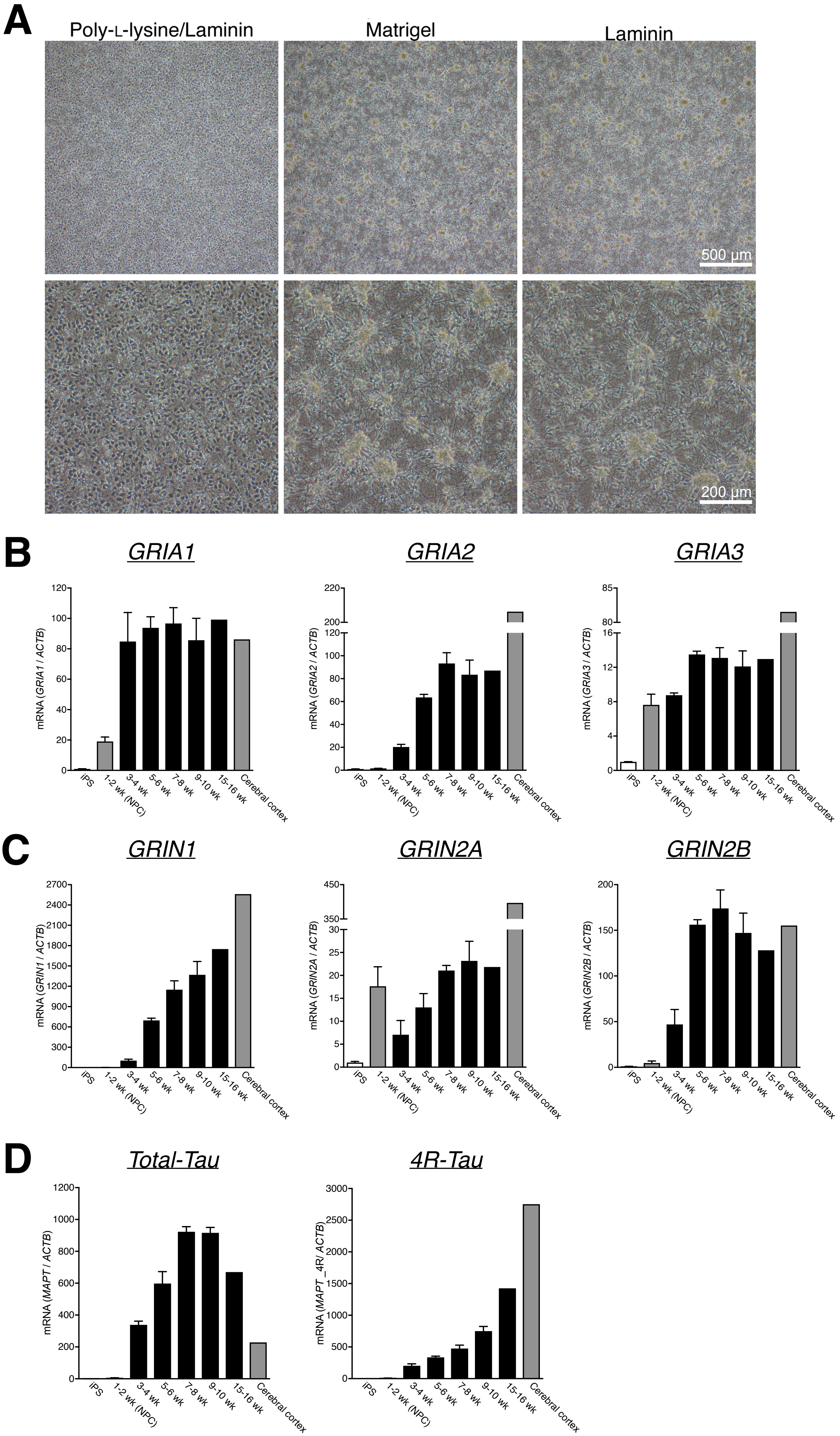


**Figure. S3 (Zhou *et al.*)**

### Sup Fig4.pdf

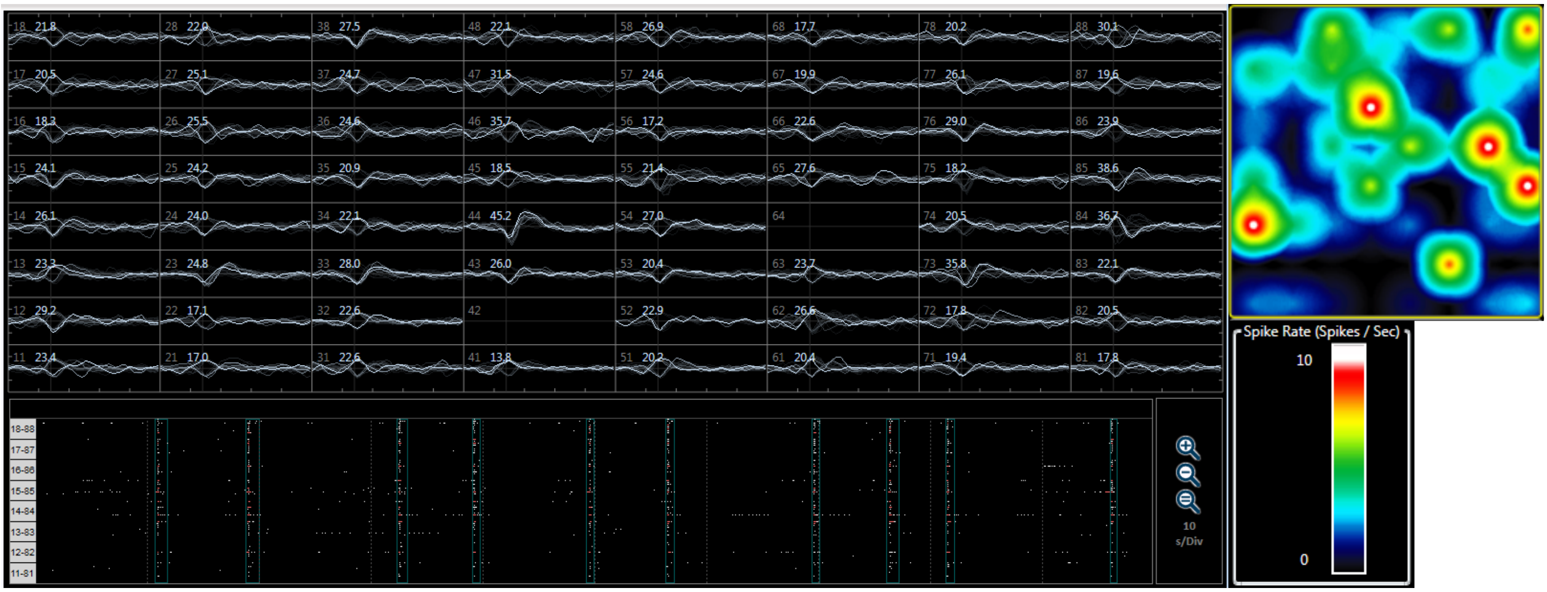


**Figure. S4 (Zhou *et al.*)**
